# A cold-water coral garden with co-occurring *Antipathella subpinnata* and *Dendrophyllia cornigera* in the mesophotic zone of the northern Bay of Biscay

**DOI:** 10.1101/2025.11.14.688465

**Authors:** Raquel Marques, Aodren Le Gal, Aymeric Boulay, Simon Fournier, Lisa Wauters, Yann Patry, Thomas Pommier, Philippe Bornens

## Abstract

Large aggregations of cold-water corals (CWC), often termed CWC gardens, are considered as biodiversity hotspots and recognized as vulnerable marine ecosystems. Their three-dimensional structure creates important habitat complexity, providing refuge, nursery and feeding areas for many deep-water species. Their vulnerability and limited current knowledge highlight the importance of describing the distribution and habitats of these gardens. Using remotely operated vehicles (ROV), we describe an unprecedented large aggregation of the co-occurring black (*Antipathella subpinnata*) and yellow (*Dendrophyllia cornigera*) coral gardens in the mesophotic in the northern Bay of Biscay. These distributions acknowledged their latitudinal and bathymetric distributions, in particular for *A. subpinnata*. Black and yellow corals presented mean densities of 2.0 ± 1.8 colonies.10 m^-2^ and 1.5 ± 1.3 colonies.10 m^-2^, respectively. Following the complex topography of this area, they were mostly fixed on hard substrates, particularly in narrow crests close to or surrounded by sand or mud planes. These findings are essential to develop appropriate local spatial management plans, ultimately contributing to the conservation of these vulnerable ecosystems.

## Introduction

Cold-Water Corals (CWC) represent a diverse group of taxa adapted to low-light, cooler environments, distinct from the shallow, sunlit habitats of typically tropical zooxanthellate corals (Freiwald et al. 2004a; Cairns 2007; Roberts and Cairns 2014). While commonly associated with the deep sea—defined as depths greater than 200 m—CWCs have also been found in habitats as shallow as 39 m (Freiwald et al. 2004a) and more typically below 50 m, within the mesophotic zone (Roberts et al. 2006; Cairns 2007). Dense aggregations of CWC often called “coral gardens”, are made up of various coral species from different taxonomic groups, such as soft corals, gorgonians and sea pens (Octocorallia), as well as black corals (Antipatharia), hard corals (Scleractinia), and, in some regions, hydrocorals (Stylasteridae) (OSPAR 2008; Roberts and Cairns 2014). CWC are amongst the most important ecosystem engineers (Jones et al. 1994) in mesophotic and deep environments, including continental shelves, slopes, canyons, seamounts and ridge systems across the globe (Roberts and Cairns 2014). These organisms form three-dimensional habitats, providing structural complexity to the ecosystems, harbouring distinct and rich communities (Freiwald et al. 2004a). CWC are slow growing, long lived, and structurally fragile organisms (Adkins et al. 2004; Sherwood and Edinger 2009; Orejas et al. 2011), which make them highly vulnerable to many anthropogenic stressors (Smith et al. 2019). Furthermore, recent genomic studies confirm the existence of weak genetic connectivity between distant populations of these organisms, highlighting the importance of isolated stepping stones populations for the long-term resilience of these species (Addamo et al. 2021; Terzin et al. 2021; Matos et al. 2024). In addition, these habitats are considered as biodiversity hotspots, providing refuge, nursery and feeding areas for many deep-water species (Baillon et al. 2012; D’Onghia et al. 2012; Henry and Roberts 2017; Soares et al. 2020). These characteristics guarantee that CWC meet nearly all criteria that characterize Vulnerable Marine Ecosystems (VME), i.e. ecosystem engineers, physically or functionally fragile, easily disturbed and with a very slow recovery capacity (FAO 2009), they are recognized by the OSPAR list of threatened and/or declining habitats (described as in “poor overall status”, OSPAR 2008) and they are considered as Habitats of Community Interest (European Union Council Directive 92/43/EEC). CWC species are listed in several conventions for the protection of marine environments, such as the annex II of the Washington convention (CITES 2024), the annex II and III of the Barcelona convention (Mediterranean) (UNEP/MAP 1995) and the annex III of the Bern Convention (CE 1979), which all aim to protect endangered or vulnerable species, though their scopes, mechanisms, and geographic coverage differ.

Among the anthropogenic stressors affecting CWC, fishing activity (especially bottom trawling) is considered to be the one causing the most serious threat, through direct physical damage or indirectly due to sediment resuspension (Ragnarsson et al. 2016; Clark et al. 2016; Yoklavich et al. 2018; Bilan et al. 2023). Likewise, the growing installation of industrial infrastructures in the marine environment, such as cables and pipes, offshore wind farms and hydrocarbon exploration platforms, or the direct seafloor exploitation, such as deep-water mining, are or may become major anthropogenic threats to the survival of these coral gardens (Kark et al. 2015; Smith et al. 2019; Simon-Lledó et al. 2019; Maxwell et al. 2022). Despite their recognised importance and vulnerability, investigations on CWC are often concentrated on deep sea areas (>200m), while mesophotic areas (ca. 50-200m) are paradoxically less well known (Angeletti and Taviani 2020, but see Loya et al. 2019), even though they represent high potential areas for the development of many of the cited anthropogenic activities. Indeed, CWC have been identified and described inside or around most submarine canyons in the Bay of Biscay (Sánchez et al. 2009; van den Beld et al. 2017; Asdal 2020; Rodríguez-Basalo et al. 2022; Abad-Uribarren et al. 2022; De Bettignies et al. 2024), but very few have been reported in the mesophotic shelf area (Freiwald et al. 2004b). Due to the vulnerability of these organisms, their functional importance, their susceptibility to increasing anthropogenic impacts, and the widespread recognition of the need for their protection, there is an urgent need to map the distribution of CWC gardens in mesophotic zones (Turner et al. 2019; Lim et al. 2021), for which fundamental knowledge is still lacking.

Anthipatarians, generally known as “black corals”, like *Antipathella subpinnata* (Ellis & Solander, 1786), and scleractinian corals, like *Dendrophyllia cornigera* (Lamarck, 1816) or “yellow coral”, are typical members of CWC assemblages.

*Antipathella subpinnata* has a white tissue and black branched skeleton, that can form large three-dimensional frameworks up to 1 m high (Bo et al. 2009). It is distributed in the Mediterranean Sea and northeast (NE) Atlantic Ocean. In the Mediterranean, the species is widely distributed and reported from 50 to 400m depth (Gori et al. 2017), with large aggregations reported in Alboran Sea,Aegean Sea (e.g. Bo and Bavestrello 2019; Rueda et al. 2021), and Italian waters (Bo et al. 2008; Chimienti et al. 2020). In the NE Atlantic this species has been found in the Azores and several seamounts, such as Josephine, Gorringe or Great Meteor (Oceana 2011; de Matos et al. 2014) and in the English Channel (Dantan 1921; Cabioc’h 1968), its northernmost report (Bo et al. 2008; de Matos et al. 2014). In the Bay of Biscay, the species was reported sporadically in several locations, such as Bay of Biscay Seamounts, in Celtic seas (Ushant island) and in the Capbreton Canyon (Grasshoff 1985; Oceana 2011; de Matos et al. 2014; Roche 2018) This species is generally found between 50 and 600 m depth, settled on hard substrates with smooth to steep inclinated seafloors and in areas characterized by strong currents (Bo et al. 2008; Oceana 2011; Wagner et al. 2012; Chimienti et al. 2020).

The yellow coral *Dendrophyllia cornigera* is a colonial scleractinian, easily identified by its bright yellow coenenchyme and irregular branching structure, creating colonies usually 30 to 40 cm tall (Enrichetti et al. 2023). This species is usually observed as scattered colonies, between 30 and 1200 m depth, with higher densities between 120 and 330 m depth, on hard or biogenic substrates on smooth to steep inclinated seafloors and soft bottoms (Gori et al. 2014; Castellan et al. 2019; Enrichetti et al. 2023). The yellow coral *D. cornigera* is present in both the Mediterranean and the northeastern Atlantic (Enrichetti et al. 2023). In the Mediterranean, this species was reported, for example, in the Alboran Sea (Ocaña et al. 2017), in the Menorca channel (Grinyó et al. 2018) in the Cassidaigne Canyon in the Gulf of Lion (Fabri et al. 2019), and in the Adriatic Sea (Mačić et al. 2024). In the Atlantic, its distribution ranges from southwestern Ireland to the Azores and the western coast of Morocco (Enrichetti et al. 2023), with rare records in the English Channel (Teissier 1965). Compared to the Mediterranean, observations of *D. cornigera* are less frequent in the Atlantic (Castellan et al. 2019). In the Bay of Biscay, this species is found from Ouessant (Girard-Descatoire et al. 1995; Castric-Fey 1996; Derrien-Courtel, et al. 2011) to Cap Breton (Ifremer 2017) and in the Cantabrian Sea (Sánchez et al. 2009; Gori et al. 2018; Rodríguez-Basalo et al. 2022), where high densities of this species have been reported (Sánchez et al. 2009; Gayá-Vilar et al. 2024). In the northern submarine canyons of the Bay of Biscay, only 3 colonies of *D. cornigera* have been reported by van den Beld et al. (2017). Despite the known presence of these two species in the Bay of Biscay, detailed information on their distribution and habitat in mesophotic shelf areas is still lacking. In addition, although the co-occurrence of *A. subpinnata* and *D. cornigera* has been previously observed in the Mediterranean (e.g. Grinyó et al. 2018) and in the NE Atlantic (Oceana 2011; de Matos et al. 2014), to our knowledge, it has never been reported in the northern Bay of Biscay. This is particularly important since both species are listed on the IUCN Red List (Mediterranean assessments): *A. subpinnata* is considered as “near threatened” (Bo et al. 2015) and *D. cornigera* as “endangered” (Orejas et al. 2015).

Here we explored an area of the mesophotic environment off the northwestern coast of France, using a Remotely Operated Vehicle (ROV) equipped with high-definition video cameras, in order to describe the distribution, density and habitat characteristics of the black (*Antipathella subpinnata*) and yellow (*Dendrophyllia cornigera*) coral gardens. This information is not only important to a better understanding of their ecology, but above all, to guide spatial management efforts and, in turn, promoting the long-term conservation of these ecosystems.

## Material and Methods

The study site was located in the northern shelf of the Bay of Biscay, in the temperate Northeast Atlantic Ocean, about 60 km offshore of the northwestern coast of France (Figure 1). It covered a total area of approximately 122 km^2^, between 70 and 95 m depth, dominated by hard and soft substrates (Figure 1). The survey area focused on an elongated rocky plate, oriented in a northwest-southeast direction. This plate was separated in two sections (northwestern and southwestern) by a mud plane of about 5 km long, which extends towards the south and southwestern area.

**Figure 1:**
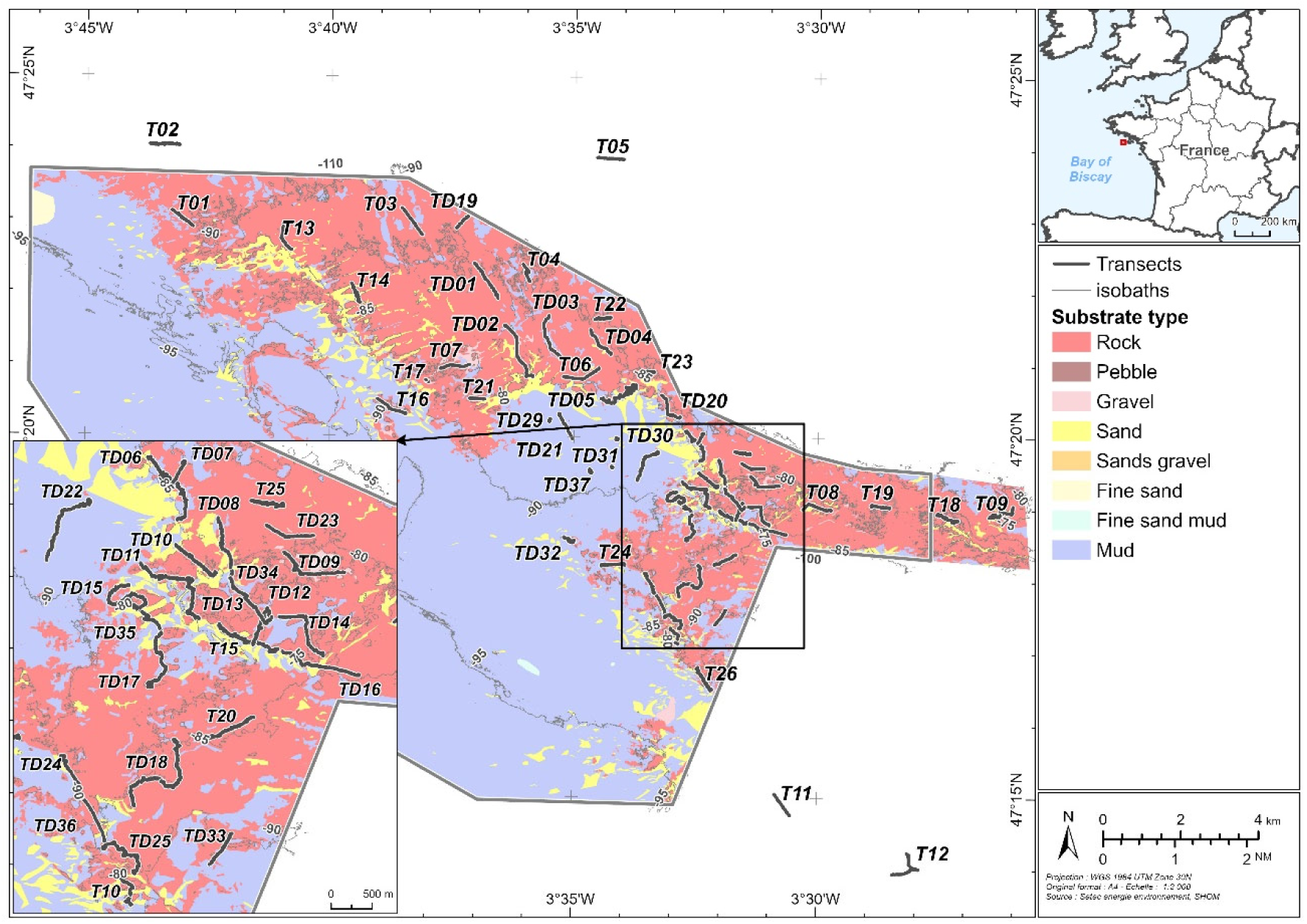
Study site at the northern Bay of Biscay, showing all ROV transects (black lines) distributed on the study area. The T transects were initially distributed in different areas of the rocky plate and complemented by the TD transects on potentially high development areas. The different substrate types of the immediate study area are shown, surrounded with unknown substrate type areas, outside of the study site (in white). The transects outside the study area (T02, T05, T11 and T12) were performed on rock substrates. Contour grey lines correspond to isobaths 80, 85, 90 and 95 m.

In total, 60 transects were conducted in the study site during four cruises (Table 1): in June 2022 (17 transects), in April 2023 (29 transects), in July 2023 (6 transects) and in September 2023 (8 transects). The transects were initially distributed in the study area in order to explore different substrate types and to characterise the benthic habitat type in the area (33 T transects). After the identification of some CWC colonies, 27 complementary transects (TD transects) were performed in areas with high potential for the development of CWC gardens, i.e. hard substrates with smooth to steep inclinated seafloors (Castellan et al. 2019) (Figure 1, Table 1).

**Table 1:**
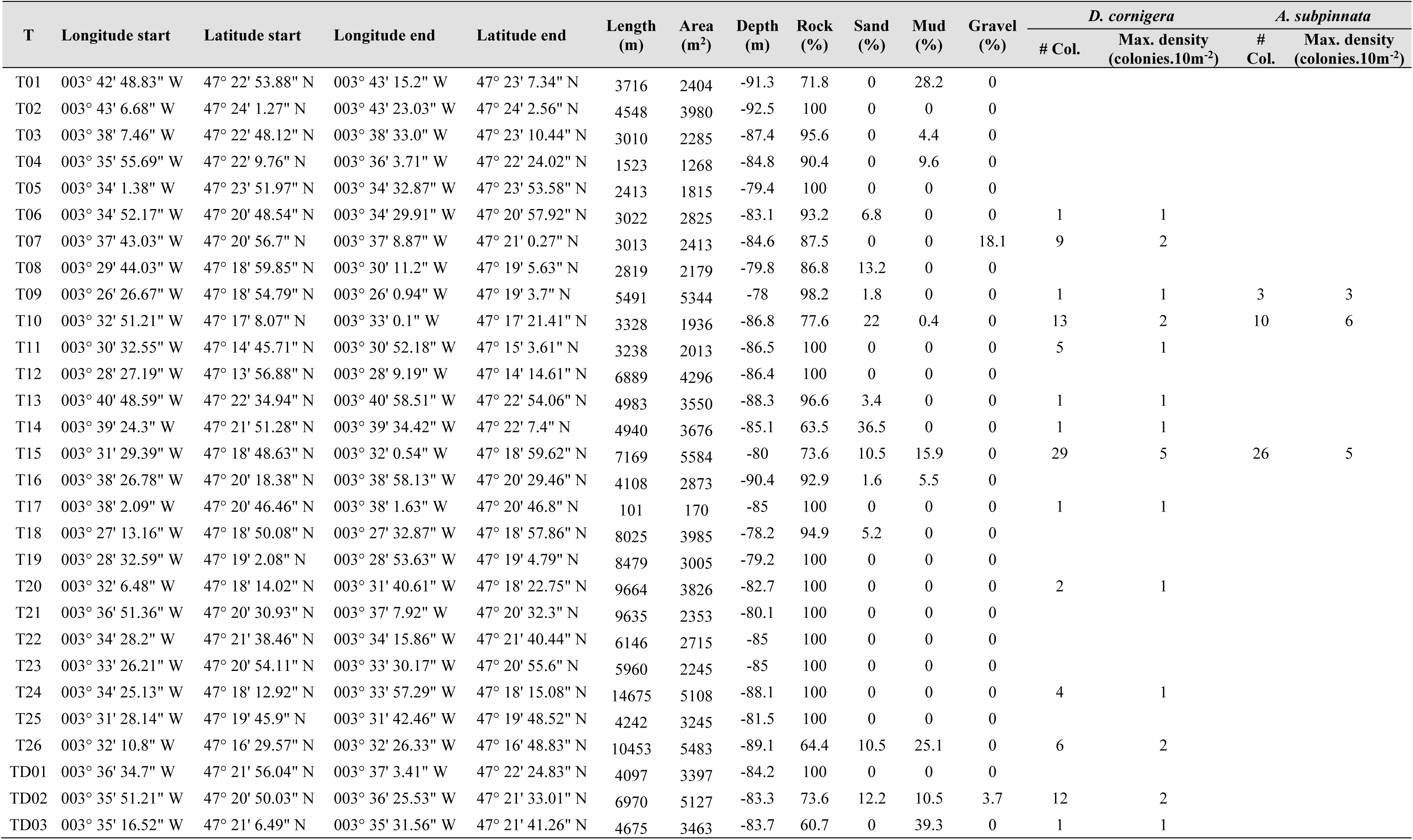
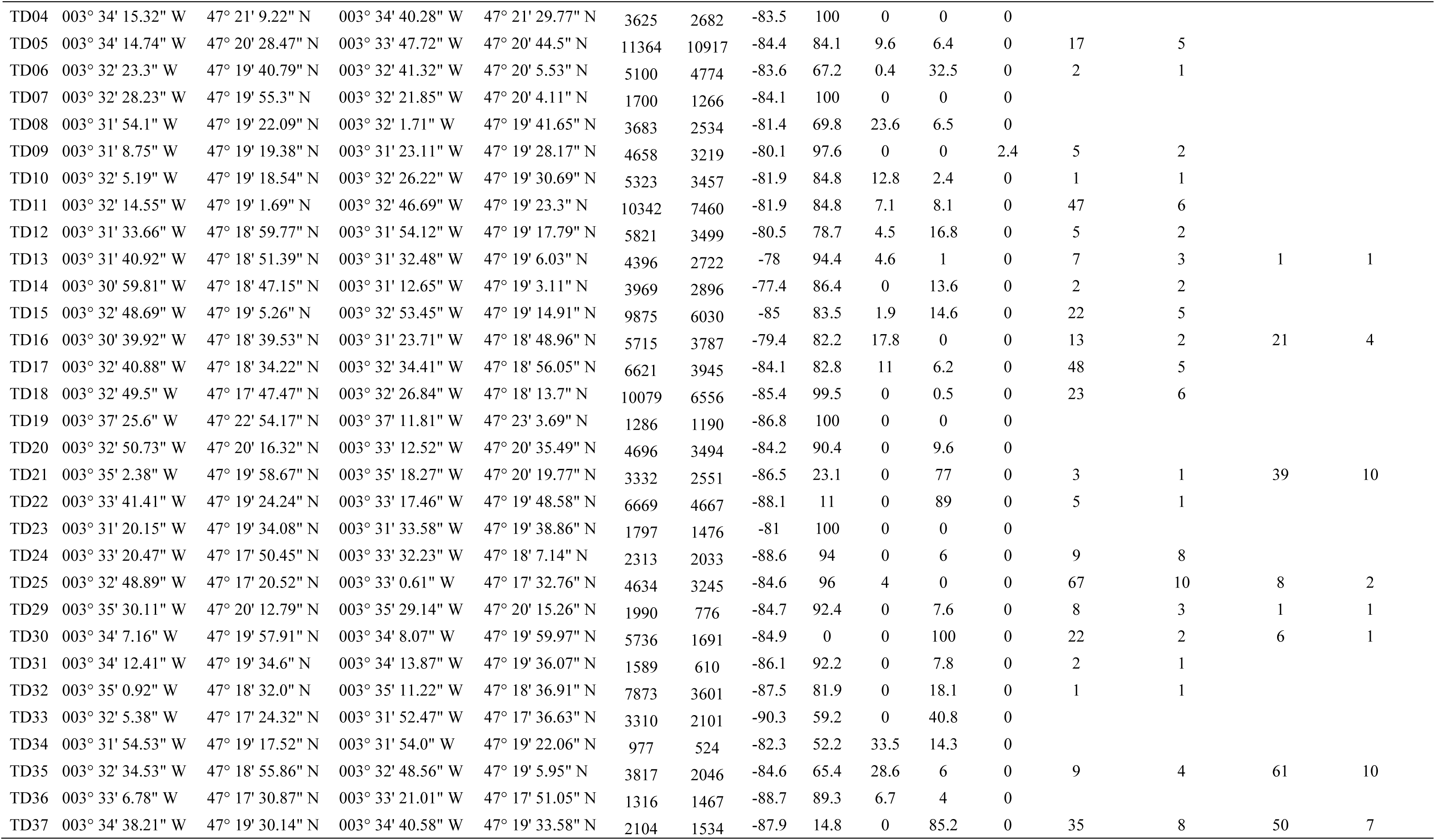
Summary information for each ROV transect: GPS coordinates of the start and end, total length and estimated surveyed area, mean depth, percentage of substrate type surveyed, total number of colonies (# Col.) and maximum density for *D. cornigera* and *A. subpinnata*.

The transects were surveyed using a Remotely Operated Vehicle (ROV, Super Achille model) equipped with two cameras: a high-definition vertical camera, that filmed the bottom at a 30° angle, and a standard definition camera with an adaptable angle, that filmed the front area of the ROV for maneuvering. The ROV was also equipped with a maximum scene illumination system of 12 000 lummens, a depth sensor, an underwater navigation system (USBL Subsea Transponder) with precise GPS location and two parallel laser beams, providing a scale of 6 cm for distance estimations during subsequent image analysis.

All transect videos were inspected using VLC multimedia player (v. 3.0.17.4 Vetinari), while the post-treatment was performed using the ArcGIS Pro and QGIS softwares (version 3.3.1 et 3.28.4, respectively). Plots were produced with R (version 4.3.2) (R Core Team 2023).

All transect videos were visually analysed by independent and well-trained experts to identify all colonies of the target species (*Dendrophyllia cornigera* and *Antipathella subpinnata*). For *D. cornigera*, accurately counting individual colonies during video analysis was sometimes challenging, as many colonies were attached on deep oyster aggregation, with some colonies not visible in the video footages. In such instances, each visible aggregation of multiple polyps was treated as a single colony, likely leading to an underestimation of the total number of colonies. When different counts occurred for the same transect, the transect was reanalysed by the three experts together. For each colony observed, depth, subtrate type and the GPS location were registered. The identification of the subtrate type of each colony was performed based on video analysis using the CATAMI classification. The CATAMI classification system (Collaborative and Automated Tools for the Analysis of Marine Imagery) is a standardized framework used to categorize and annotate benthic marine organisms and the physical habitat characteristics from underwater imagery (Althaus et al. 2015). The physical component of the CATAMI classification is based on three main characteristics: substrate, relief and bedform (Althaus et al. 2015). The CATAMI substrate categories found in the study area were consolidated rock, consolidated boulders and sand / mud.

The bathymetry (resolution of 2 x 2m) and the substrate type of the whole study site were obtained from SHOM (https://data.shom.fr). The substrate type from SHOM was used for background presentation of the distribution of substrate types in the study area (Figure 1) and to estimate the percentage of each substrate type surveyed during the ROV transects (Table I), calculated as the percentage of the ROV transect on each substrate type, according to their geographic position. A slope map was calculated from the bathymetric data using the function “slope” in QGIS and the associated slope value of each colony was then obtained according to their geographic position.

The surface area explored in each transect was estimated as following: For each transect, 10 random printscreens of the videos (total of 570) were used to calculate an average width of the field of view using the laser scale (width = 1.65 m ± 0.68 m). The distance of the ROV to the bottom was maintained as consistently as possible during video aquisition because of low visibility, so the field of view was assumed to remain constant across all transects. The length of the transects ranged between 100 and 14675 m long, depending on the benthic topography. The estimated surface area was calculated by multiplying the field of view by the length of the transects, excluding the superposed surfaces where the ROV transects intersected, during the post-treatment. The estimated surface area examined per transect varied between 170 to 10917 m^2^ (Table 1). The transects were then split in sections of 10 m^2^ to calculate colony density (colonies.10m^-2^). The surface area occupied by each species was estimated by summing all 10 m^2^ section where colonies were observed.

## Results

A total of 191 308.8 m^2^ were inspected in the study area where a total of 665 colonies of CWC were observed: 439 colonies of *D. cornigera* and 226 colonies of *A. subpinnata* (Figure 2, Table 1), covering a total estimated area of 2760 and 1020 m^2^, respectively (0.5 and 1.4% of the inspected area, respetively). Most colonies (97%) were located around the mud plane that separates the rocky plate in two sections, particularly in the southeastern area (Figure 2). *D. cornigera* were more widely distributed in the area, being observed in 37 different transects (1 to 67 colonies per transect), while *A. subpinnata* were only observed in 10 transects, with up to 50 colonies observed per transect (Table 1). The colonies of *A. subpinnata* were in most cases observed in areas where the colonies *D. cornigera* were also present, except in some sections of the transects T09, TD16 and TD21 (Figure 2). Both species of CWC were absent in 23 out of 60 transects, which were mainly distributed in the most northern areas of the whole surveyed rocky plate, away from the southern large mud plane (Figure 1 and 2).

**Figure 2:**
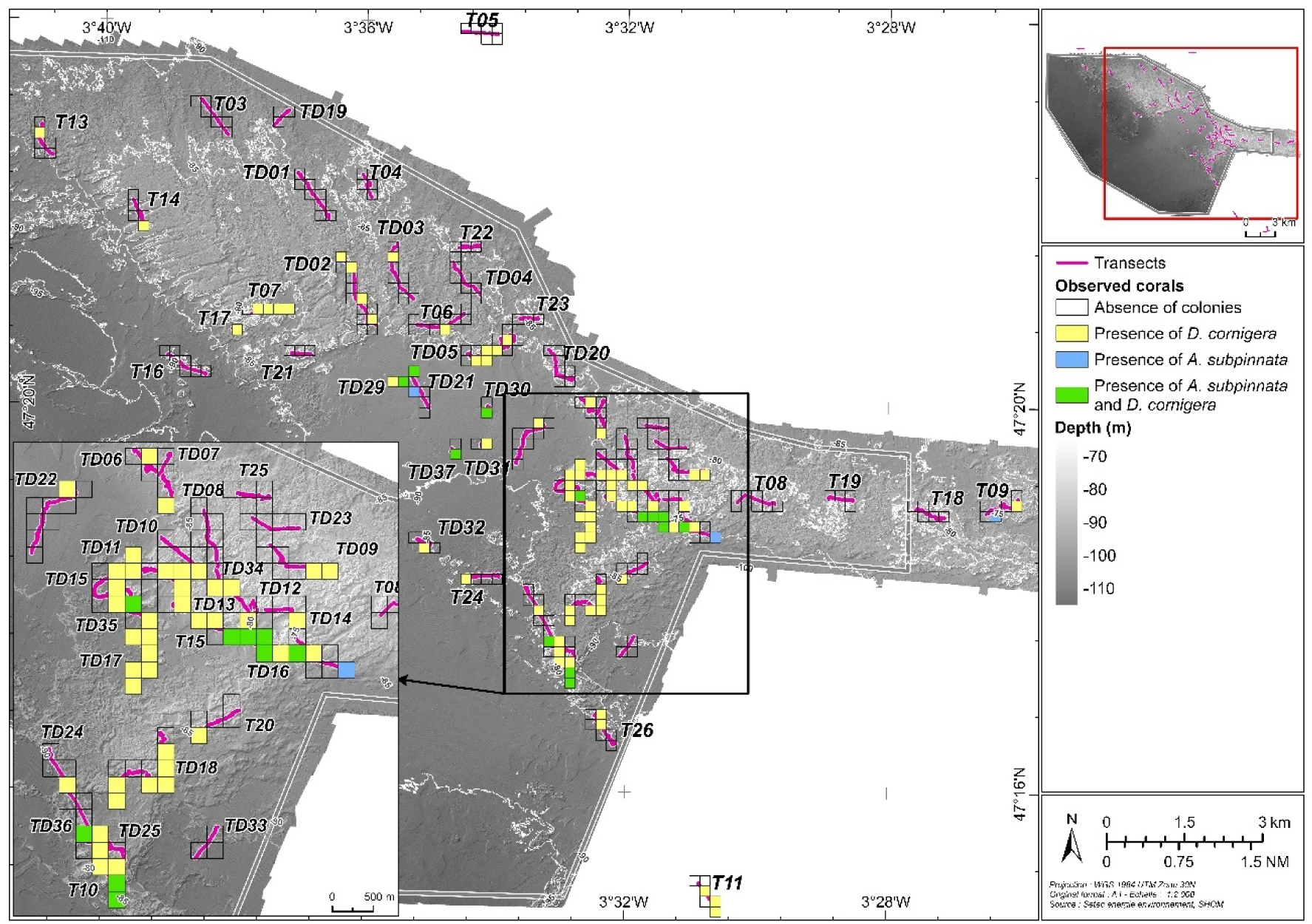
Distribution of the CWC colonies in the study area. Each square represents an area of 250 m^2^, indicating the (co-)occurrence of *A. subpinnata* and *D. cornigera*. No colonies were observed in the transects T01, T02 and T12 that are out of the map range.

The density of CWC in the study site ranged 1 to 10 colonies.10 m^-2^ for both species with an average density (when present) of 2.0 ± 1.8 colonies.10 m^-2^ and 1.5 ± 1.3 colonies.10 m^-2^, for *A. subpinnata* and *D. cornigera*, respectively. The transects with high densities of both co-occurring species were T15, TD25, TD35 and TD37 (Table 1) detailed in Figure 3. The transects with maximum densities of *D. cornigera* were TD25 (10 colonies.10 m^-2^), TD24 (8 colonies.10 m^-^2) and TD37 (8 colonies.10 m^-2^). Maximum densities of *A. subpinnata* were observed in transects TD21, TD35 and TD37, where values of 10, 10 and 7 colonies.10 m^-2^ were respectively registered (Figure 2 and Figure 3). The areas with high densities of CWC were often characterised by rocky formations or outcrops, with pronounced reliefs, forming submarine rocky wall edges, and frequently close to or even isolated in areas of sand or mud planes. The colonies were typically settled on top of those cliffs or on the edge of steep walls (Figure 3). Although the size of colonies was not consistently measured, the visual inspection of videos revealed that the colonies of *D. cornigera* ranged from a small size (few polyps) to big colonies of more than 30 different polyps, while the size of *A. subpinnata* was highly variable, with some colonies reaching more than 1m high. A video with extractions of some transects is provided in supplementary material.

**Figure 3:**
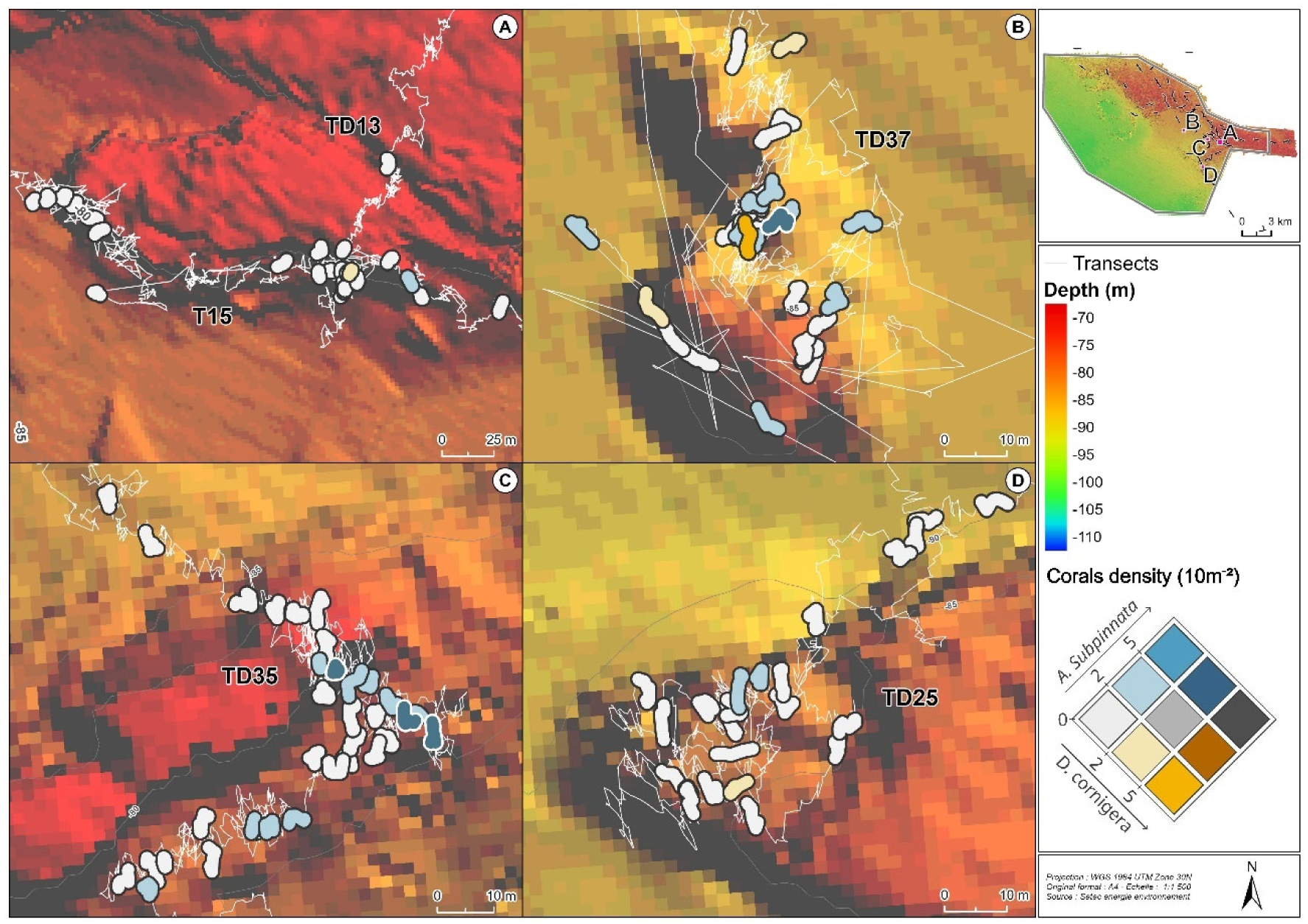
Density of CWC in transects with high densities of co-occurring *D. cornigera* and *A. subpinnata*. The density of colonies is presented in a bivariate scale: present but with low densities in white, high densities of only *D. cornigera* in orange, high densities of only *A. subpinnata* in blue and high densities of both species in dark grey. White lines represent the ROV transects. Each polygon represents sections of the transect of 10m^2^. Their shape is influenced by the complexity of the ROV transects: a linear transect section is represented by longer polygons, while twisted sections are represented by rounded and apparently smaller polygons.

*A. subpinnata* were all observed directly fixed on rock substrates, often covered with a thin layer of sediment. Indeed, according to CATAMI classification, 98.7% of the observed colonies were fixed on consolidated rock, while only 3 colonies (1.3%) were fixed on consolidated boulders (Figure 4b). Likewise, most *D. cornigera* colonies (89.9%) were also fixed on consolidated rock substrates, while 10.1% were fixed on consolidated boulders (Figure 4b). In some cases, rock substrate was covered by oysters (*Neopycnodonte cochlear*) reef that create nodules used then as settlement substrate for this species. All colonies of *A. subpinnata* and *D. cornigera* were observed between 75 and 91m depth (Figure 4b). Higher occurrences of *A. subpinnata* were observed at 83 and 87m depth while *D. cornigera* occurred mainly around 85m depth. For both species, most colonies were observed on flatter areas, with 58 and 62% of the colonies of *A. subpinnata* and *D. cornigera*, respectively, fixed on substrates with slopes less than 10° (Figure 4d). In areas with steeper slopes (> 35°), *A. subpinnata* was more frequent (11% of all colonies), than *D. cornigera* (5.5% of all colonies).

**Figure 4:**
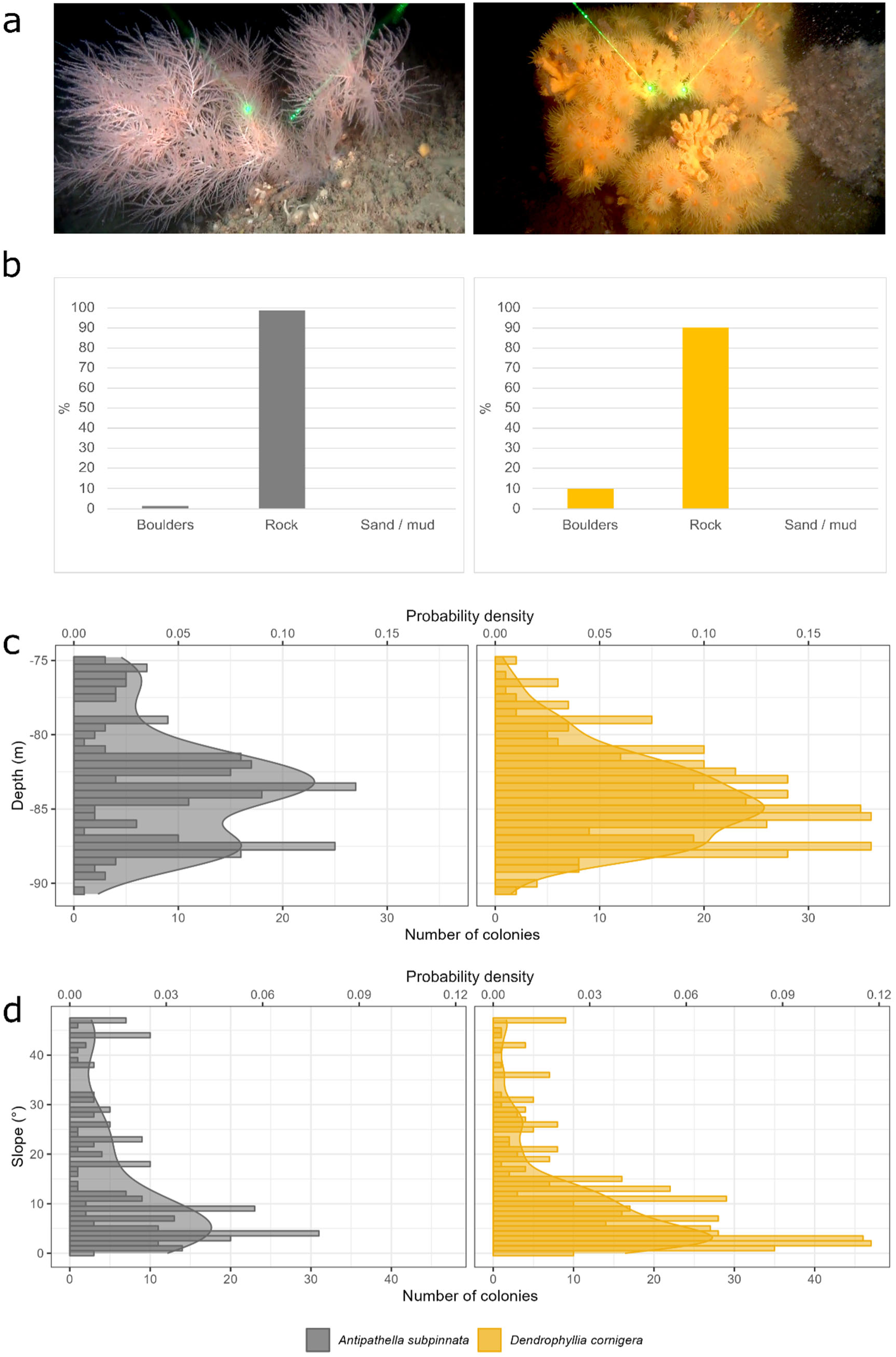
Description of habitat of *A. subpinnata* (left) and *D. cornigera* (right). a) ROV images of both species (distance between laser beams of 6 cm), b) settlement substrate type, according to CATAMI classification system, and histogram (bars) and probability density (curves) of colonies by depth (c) and bathymetric slope (d).

## Discussion

Despite the recognized distribution of different species of CWC in the submarine canyons of the Bay of Biscay, little is known on the distribution, density, and habitat type of these organisms in the shallower mesophotic areas. Here we describe the biggest CWC garden of two co-occurring species, *A. subpinnata* and *D. cornigera,* in the northern shelf area of the Bay of Biscay ever reported so far. While both species have been previously reported in the NE Atlantic, this study presents large populations observed near their most upper bathymetric and northern latitudinal distribution limit, in particular for *A. subpinnata*.

The extensive sampling effort in this study allowed to find and describe a population of *A. subpinnata* and *D. cornigera*, covering, a total estimated area of 3,610 m^2^ (cumulative area of 10m² sections with at least one coral colony), with maximum densities of 10 colonies.10 m^-2^ (*i.e.* 1 colony.m^-2^) for both species. For *A. subpinnata*, its average density of 2.0 ± 1.8 colonies.10 m^-2^ (0.20 ± 0.18 colonies.m^-2^) was lower than the only reported densities of these species in the Atlantic, the *A. subpinnata* garden found in deep waters (> 150m) off the Azores, with an average and maximum density of 0.75 ± 0.72 and 2.64 colonies.m^-2^, respectively (de Matos et al. 2014). The density of *A. subpinnata* was also lower than the largest populations found in the Mediterranean Sea, such as in Southern Tyrrhenian Sea (average of 1.4 and up to 5.2 colonies m^-2^) (Bo et al. 2009), or in Tremiti Islands (average between 0.22 ± 0.03 and 2.40 ± 0.26 and up to 7.2 colonies m^-2^) (Chimienti et al. 2020). However, similar densities were reported in areas with important gardens at over 100m depth, such as in Northern Tyrrhenian Sea (average of 0.27 ± 0.10 and 0.13 ± 0.04 colonies m^-2^) (Bo et al. 2014) and in the Sardinia Channel (0.11 ± 0.06 colonies m^-2^) (Cau et al. 2015). For *D. cornigera*, its average density of 1.5 ± 1.3 colonies.10 m^-2^ (0.15 ± 0.13 colonies.m^-2^) was consistent with 90% of the reports in Italian waters (< 1.6 colonies m^-2^) but much lower than the maximum values reported in this area (48 colonies m^-2^) (Enrichetti et al. 2023). In the southern Bay of Biscay, recent studies observed this species at high average densities of 3.1 to 5.0 colonies.m^-2^, with sporadic peaks of densities of 60.6 colonies.m^-2^ (Gayá-Vilar et al. 2024) estimated on the basis of under 2m² images. However, we highlight that the densities presented in this study are likely underestimated: firstly, we calculated densities per 10 m² sections, which may have overlooked finer density variability within each section. It is, therefore, possible that higher densities were present at smaller spatial scales; secondly, when *D. cornigera* colonies were fixed on boulders of oyster nodules, it was visually difficult to distinguish individual colonies, so they were often counted as a single colony, further underestimating density for this species. Considering these biases, and despite the lower densities of these two species in comparison with other CWC gardens, their presence in an extensive area in mesophotic environment is remarkable.

The populations of *A. subpinnata* and *D. cornigera* reported in this study were observed at depths between 75 and 92m, with peak occurrences around 85m. *A. subpinnata* inhabit depths lower than 100m, but the majority of populations occur between 200 and 600 m depth (Bo et al. 2009). *While* Bo et al. (2008) classified A. subpinnata as stenothermal based on its bathymetric range (<15°C), Godefroid et al. (2023) demonstrated short-term thermal resilience up to 19°C in controlled ex situ conditions. In the Bay of Biscay, colonies of *A. subpinnata* have been already reported off the coast of Aber Wrac’h (Finistère) at -54 meters (Dantan 1921), but large populations in mesophotic habitats as in this study have never been described. In contrast, the higher thermal tolerance of *D. cornigera* (Reynaud et al. 2021) allows its broad bathymetrical distribution from 30 meters (exceptional upwelling site) to 1,200 meters (Castric-Fey 1996; Castellan et al. 2019; Enrichetti et al. 2023), but its preferential bathymetric distribution in the Atlantic seems to be between 100 and 200 meters (Castellan et al. 2019). In the northern Bay of Biscay, some mesophotic observations have been reported in the bays of Douarnenez and Audierne (Toulemont 1972), in the Penmarc’h area (Doré 2012), in the Glénan sector (Girard-Descatoire et al. 1996), as well as off Belle-Ile (Castric-Fey 2001). However, these are isolated observations with low or unknown densities of individuals.

The colonies of *A. subpinnata* and *D. cornigera* are typically observed on rock substrates, such as isolated rocky outcrops, cliff walls and rocky plateaus (e.g. Bo et al. 2008, 2009; Castellan et al. 2019; Enrichetti et al. 2023). Likewise, in this study, the areas with higher densities of these species were often characterised by rocky formations with pronounced reliefs, where these corals seem to favor local topographic highs, frequently close to sand or mud planes or even fixed on isolated outcrops in these planes. The submarine basin of mud plane of 5 km long, surrounded by topographic highs, appears to favor the development of CWC gardens in the area. These results indicate that, despite the limited sporadic reports of these species in the Bay of Biscay, the area offers suitable conditions for the development of large populations (Hall-Spencer et al. 2007; Reveillaud et al. 2008). Indeed, CWC occur in areas where topography and currents generate an accelerated flow (Rogers 1990; De Mol et al. 2002; Freiwald et al. 2004a; Reveillaud et al. 2008). Internal tidal waves, which are strong in the shelf area of the Bay of Biscay (Lazure et al. 2009), are responsible for the vertical and bottom mixing, that further enhance coral food supply through organic matter resuspension (Frederiksen et al. 1992; Lesser et al. 2009). Nutrient-rich waters boost phyto- and zooplankton productivity, providing a major food source for corals (Freiwald et al. 2004a; Gori et al. 2015), supporting higher fatty acid contents and improving CWC fitness (Büscher et al. 2017; Gori et al. 2018). The resistant chitinous skeleton of *A. subpinnata* and the scleractinian skeleton of *D. cornigera* are particularly suitable for living in strongly moving water (Reveillaud et al. 2008; Bo et al. 2009; Chimienti et al. 2020). Indeed, currents rich in suspended matter are probably the major environmental constraints influencing the development of CWC gardens (Castric-Fey 1996; Bo et al. 2009; de Matos et al. 2014; Chimienti et al. 2020). The strong and complex bottom and slope currents of the Bay of Biscay, influenced by wind, tides, freshwater input from large rivers and by the strong continental slope with numerous canyons (Batifoulier et al. 2012; Charria et al. 2013; Porter et al. 2016; Borja et al. 2019), can therefore provide the suitable hydrodynamic conditions for these species. The presence of most colonies at the edge of the cliffs and on steeper inclinations (particularly *A. subpinnata*) are likely associated with fine scale bottom current patterns. We suspect that these currents contribute to clear sediment deposition, to supply resuspended organic matter from the southern mud plane to the suspension feeding corals, to provide a sufficiently rapid renewal of water needed for respiration, while the steeper slopes avoid the accumulation of detritus (Castric-Fey 1996; Bo et al. 2009, 2011; Chimienti et al. 2020).

The (co)occurrence of these species in the northern Bay of Biscay seems to represent, to our knowledge, a geographically isolated CWC garden, particularly for *A. subpinnata.* This is particularly important since isolated mesophotic coral populations are recognized as essential for vertical and horizontal connectivity (reviewed in Soares et al. 2020). Recent genomic studies on *A. subpinnata* from the Mediterranean Sea confirm the existence of weak genetic connectivity between offshore (>150m depth) and coastal populations (65-75m depth) (Terzin et al., 2021), highlighting the role of isolated populations in mesophotic areas, as stepping stones to increase genetic connectivity through larval dispersal at least in this area. This underscores the importance of documenting the presence of *A. subpinnata* in this unique site. Such a discovery expands our understanding of the species’ distribution in the Northeast Atlantic and highlights a potentially isolated but ecologically significant habitat. Given the rarity of observations, this population could serve as a key reference point to enhance knowledge of the species’ habitats and connectivity dynamics. Furthermore, these species are sensitive to anthropogenic disturbances, particularly fishing activities (Clark et al. 2016; Orejas et al. 2019; Bilan et al. 2023), and industrial infrastructures (Smith et al. 2019). Indeed, we occasionally observed ghost fishing lines entangled on *D. cornigera* colonies, underlining the need to describe the presence and habitat type of CWC gardens in the northern Bay of Biscay (Lim et al. 2021).

**The findings of this study provide critical knowledge for the future spatial management of this area—including fisheries and marine energy planning—and ultimately support the conservation of these ecosystems.**

## Acknowledgment

First of all, the authors would like to thank to the whole biodiversity team of the *Setec energie environnement – setecinvivo* for their exceptional contributions during sampling and video analysis for this study. We would also like to thank the expertise and contribution of the SAAS (Ship as a Service) team and, particularly, the ROV pilots and the Minibex ship team. Finally, we are grateful to Luc Cadot for his careful proofreading and language corrections.

## Ethical approval

No animals were collected or harmed for this article and did not require ethical approval for research.

## Contributions

YP, ALG and PB conceived the study. ALG performed material preparation, data collection and was the leading expert of video analysis. SF and LW analysed the videos. RM and AB performed the data analyses. RM, ALG and TP wrote the manuscript. All authors revised and approved all versions of the manuscript.

